# Oleocanthal and oleocanthal-rich olive oils induce lysosomal membrane permeabilization in cancer cells

**DOI:** 10.1101/610097

**Authors:** Limor Goren, George Zhang, Susmita Kaushik, Paul Breslin, Yi-Chieh Nancy Du, David A. Foster

## Abstract

Oleocanthal is a phenolic compound found in varying concentrations in extra virgin olive oil Oleocanthal has been shown to be active physiologically, benefiting several diseased states by conferring anti-inflammatory and neuroprotective benefits. Recently, we and other groups have demonstrated its specific and selective toxicity toward cancer cells; however, the mechanism leading to cancer cell death is still disputed. The current study demonstrates that oleocanthal, as well as naturally oleocanthal-rich extra virgin olive oils, induced damage to cancer cells’ lysosomes leading to cellular toxicity *in vitro* and *in vivo*. Lysosomal membrane permeabilization following oleocanthal treatment in various cell lines was assayed via three complementary methods. Additionally, we found oleocanthal treatment reduced tumor burden and extended lifespan of mice engineered to develop pancreatic neuroendocrine tumors. Finally, following-up on numerous correlative studies demonstrating consumption of olive oil reduces cancer incidence and morbidity, we observed that extra virgin olive oils naturally rich in oleocanthal sharply reduced cancer cell viability and induced lysosomal membrane permeabilization while oleocanthal-poor oils did not. Our results are especially encouraging since tumor cells often have larger and more numerous lysosomes, making them especially vulnerable to lysosomotropic agents such as oleocanthal.

## Introduction

Olive oil has been consumed by humans for millennia and is frequently associated with health-related properties. The Mediterranean diet and olive oil consumption in particular are correlated with lower cancer incidence and mortality (1–3). A meta analyses of nineteen observational studies performed between 1990 and 2011 that included approximately 35,000 individuals found that olive oil intake is inversely related to cancer prevalence (4). More recently, a randomized trial found that women who adhered to a Mediterranean diet supplemented with extra virgin olive oil (EVOO) had 62% less invasive breast cancer incidence than a control group that was advised to restrict dietary fats (5). These studies, however, did not distinguish between the protective effects of EVOO’s triglycerides and its phenolic components. Furthermore, to the best of our knowledge, no controlled study tested the effect of high phenolic EVOO on cancer.

(-)-Oleocanthal, also known as deacetoxy-ligstroside aglycon, was first identified as a minor phenolic compound in the fruit of the olive tree by Motedoro et al. in 1993 (6). A few years later, it was reported to be the primary agent that conveys strong stinging sensation at the back of the throat when ingesting certain EVOOs (7). In 2005, Beauchamp and colleagues published the first paper to refer to this compound as oleocanthal [Oleo - for oil, Canth – Greek for stinging or literally prickly (named for the throat irritation caused by oleocanthal), and Al – for the two aldehyde groups that are believed to be responsible for oleocanthal’s reactivity] (8). Beauchamp and colleagues also identified oleocanthal as a potent inhibitor of cyclooxygenase enzymes, conferring anti-inflammatory activity that was more potent than ibuprofen (8). Since this pioneering oleocanthal paper was published, several groups looked at its medicinal neuroprotective properties. Pitt et al. observed that low doses of oleocanthal altered Alzheimer’s-associated amyloid-β oligomers (9), and Li et al reported that oleocanthal inhibited tau fibrillization (10). In 2018, Batarseh et al. reported that high-oleocanthal EVOO reduced amyloid-β load and related toxicity in a mouse model of Alzheimer’s disease. (11)

Since oleocanthal inhibits cyclooxygenase enzymes that mediate inflammation, which is associated with cancer initiation and progression, there was increasing interest in studying the anti-cancer properties of Oleocanthal. The first reports of oleocanthal attenuating tumorigenicity were in 2011. Elnagar et al. (12) demonstrated oleocanthal’s ability to prevent cell migration in a metastatic model for the breast carcinoma cell line MDA-MB-231. Khanal et al.(13) showed that oleocanthal inhibits phorbol ester-induced cell transformation in murine JB6 Cl41 cells. Additionally, the authors showed that oleocanthal inhibits proliferation and the ability to form colonies formation in soft agar of HT-29 colon cancer cells. Other reports illustrated the anti-proliferative activity of oleocanthal using various cancer cell lines – including breast carcinoma (14), multiple myeloma (15), hepatocellular carcinoma (16), melanoma (17), and prostate cancer (12). The proposed mechanisms for oleocanthal activity differ among the different research groups. El Sayed and colleagues published several papers illustrating that oleocanthal acts a c-Met inhibitor (12, 14, 18). c-Met is a receptor tyrosine kinase that is activated by hepatocyte growth factor. c-Met acts upstream of both the PI3K and the MAPK pathways. The El Sayed group demonstrated oleocanthal inhibitory effects on c-Met downstream targets when cells were stimulated with hepatocyte growth factor. In their studies, the mode of death was apoptotic as evident by the induction of cleaved caspase-3 and cleaved PARP. Pei et al. reported that oleocanthal inhibits growth and metastasis of hepatocellular carcinoma by blocking activation of the transcription factor STAT3 (16). Interestingly, one of the canonical ways in which STAT3 is activated is via receptor tyrosine kinases, including c-Met. Yet the upstream initiating event that promotes cancer cell death remains uncertain.

Whereas we previously reported that oleocanthal induces cancer cell death via lysosomal membrane permeabilization (LMP) (19), other mechanisms for cancer cell death – notably apoptosis – in response to oleocanthal have been reported (16, 20, 21). Different forms of stress can induce LMP, which causes release of intra-lysosomal enzymes to the cytoplasm – resulting in lysosome-dependent cell death (22). This newly appreciated cell-death mechanism is gaining interest, since transformed cells are often characterized by a large increase to their lysosomal compartment and are strongly dependent on lysosomal function (23). Lysosomes contain over 50 different hydrolases, and many of these are up-regulated and utilized by cancer cells, often in secreted forms, for purposes of invasion, angiogenesis, and progression (24, 25). The increased reliance on lysosomal processes might also represent an Achilles’ heel for cancer. As Christian DeDuve noted – the high concentration of degradative enzymes in lysosomes make them in essence “suicide bags” (26). Lysosomes in transformed cells are more susceptible to rupture, causing release of hydrolases such as cathepsin (generic name for lysosomal proteases) into the cytosol (27). Depending upon the degree of LMP, both apoptotic and non-apoptotic death can be observed (22, 28). Low levels of LMP injures cells and triggers apoptotic death mechanisms, whereas high levels of LMP kills cells rapidly and directly as a form of necrosis.

In this report, we demonstrate oleocanthal’s ability to induce severe LMP in a variety of cancer cells lines, leading to rapid necrotic cell death *in vitro* and shrinkage of tumors and extension of lifespan in an *in vivo* mouse model for pancreatic neuroendocrine tumors (PanNET). Strikingly, we were also able to replicate the beneficial effects of purified oleocanthal by treating cells with EVOOs that naturally contain high levels of oleocanthal.

## Materials and methods

### Reagents

Oleocanthal extracted from EVOO was obtained from Dr. Alexios-Leandros Skaltsounis at the University of Athens, Department of Pharmacology. The structure and purity (97%) of the oleocanthal was determined by HPLC and H1 NMR analysis. The Governor premium EVOO limited edition (Corfu, Greece) and Atsas EVOO (Cyprus) were a gift from the producers. California Olive Ranch^TM^ EVOO (California, USA), Colavita mild olive oil (Italy), Colavita EVOO (Italy), and Mazola corn oil (USA) were purchased at a New York City grocery store. All treatments used EVOO from newly opened bottles that were kept in the dark at room temperature within one month of opening. Oleocanthal concentration was determined by H1 NMR analysis by a third party (Numega Labs, San Diego, California). All other reagents, unless noted otherwise, were purchased from Fisher Scientific.

### Cells and cell culture conditions

PC3, MDA-MB-231, MCF7, HEK-293T, MCF10A, and BJ-hTert cells used in this study were obtained from the American Type Tissue Culture Collection. Mouse PanNET N134 cells were generated by the Du laboratory(29). PC3 cells were maintained in F-12K medium, MCF10A cells were maintained in MEGM Mammary Epithelial Cell Growth Medium Bullet Kit (Lonza) supplemented with 100 ng/ml cholera toxin. other cells were maintained in Dulbecco’s Modified Eagle Medium (DMEM), supplemented with 10%, or 15% (N134) fetal bovine serum (Hyclone). No further authentication was performed.

### Antibodies

Mouse anti human galectin-3 antibody (BD Bioscineces, 556904), goat anti-human Cathepsin B antibody (R&D systems AF953), goat anti human cathepsin-D antibody (Santa Cruz sc-6486), goat anti mouse Cathepsin L antibody (R&D systems AF1515), mouse-anti human LAMP2 antibody (abcam 25631), rat anti-mouse Lamp2 antibody (Hybridoma bank 1B4D), rabbit anti-GAPDH antibody (Cell signaling 2118S), rabbit anti-HSP70 antibody (Proteintech 10995).

### Cell viability

(2,3-bis-(2-methoxy-4-nitro-5-sulfophenyl)-2H-tetrazolium-5-carboxanilide) (XTT) reduction assay was used to measure cells viability. In brief, 5×104 cells/500 μl/well were seeded into 24-well plates in triplicates. After 24 hours, cells were given treatment medium containing 20 μM oleocanthal, or vehicle only and incubated at 37°C with 5% CO2. After a 24 h incubation period, cells were treated with 150 μl XTT (Invitrogen™ Molecular Probes™ XTT cat. no. x6493) for 2 h. Then, plates were read at 480 nm wavelength by a spectrophotometer (Molecular devices, SpectraMax i3). After subtracting blank well absorbance, the absorbance of vehicle treated cells was set to 100%, and the relative absorbance of oleocanthal treated cells was reported as % viable cells.

### Lentiviral-based overexpression of HSP70

PC3 cells were transduced with either HSP70-1 (Santa Cruz biotechnology sc-418088-LAC) or control (Santa Cruz biotechnology sc-437282) lentiviral CRISPR activation particles per manufacturer protocol. Stable cell lines of HSP70 overexpressing and mock transduced control cells were generated via antibiotic selection. Viability assay was performed as described above.

### β-hexosaminidase latency assay

To determine possible direct effects of oleocanthal on lysosome stability, we examined β-hexosaminidase release from lysosomes. Briefly, fractions enriched in lysosomes were incubated with oleocanthal. After incubation lysosomes were separated from the incubating media by filtration through a 96-well plate with 0.22 μm filter using a vacuum manifold. β-hexosaminidase activity in the media was measured using a colorimetric assay as described previously (30). Broken lysosomes were calculated as the percentage of total lysosomal hexosaminidase activity detected in the flow-through.

### NMR

Oleocanthal content in oil was assessed via H-1 NMR as previously described (31). Briefly, oil samples (240 ± 20 mg) and Syringaldehyde internal standard were dissolved in 0.6 ml of CDC13. H1 NMR experiments (NS=512) were recorded on Bruker AV500. Proton signals of aldehydes from oleocanthal (9.18 ppm) and Syringaldehyde (9.77 ppm) were integrated.

### Apoptosis / Necrosis assay

Mode of death was detected by flow cytometric analysis of annexin V-FITC and propidium Iodide staining (Vibrant apoptosis assay), Molecular Probes V-13242) per manufacturer’s protocol.

### Immunohistochemistry

The Aits, Jaattela, and Nylandsted protocol for detection of damaged lysosomes by Galectin-3 translocation was performed as previously described (32, 33). Slides were visualized on confocal microscope (Nikon Instruments A1 Confocal Laser Microscope Series equipped with NIS-Elements acquisition Software).

### LysoTracker assay

2.5×10^5^ cells per well were grown in a 6 well plates. The next day, the media was changed and cells were incubated with treatment media containing 20 μM oleocanthal, 2 mM LLOMe, or DMSO for the indicated amounts of time. In the last 15 minutes of the treatment, 50 nM LysoTracker green (Invitrogen™ Molecular Probes™ LysoTracker™ green DND-26 L7526) was added to the media. Cells were harvested with trypsin EDTA, and re-suspended to 1 × 10^6^ cells/ml. Green fluorescent intensity was immediately analyzed by flow-cytometry (Orflo MoxiGo II).

### Cell fractionation and western blot analysis

Cytosolic and light membrane fractions containing lysosomes were obtained using a cell fractionation kit (Abcam ab109719) and procedure was carried according to manufacturer’s protocol. Where indicated, highly purified lysosome enriched fractions were isolated through centrifugation in discontinuous gradients of metrizamide and Percoll as previously described. Cytosolic and light membrane fractions were obtained and protein concentration was estimated. Twenty micrograms of proteins were loaded into wells of freshly prepared polyacrylamide gel. Proteins were electrophoresed and transferred to a nitrocellulose membrane. The membranes were blocked in 5% milk in PBST and incubated overnight with indicated antibodies. The membranes were washed and incubated with the appropriate secondary antibodies for one hour at RT, washed again and visualized using KwikQuant^TM^ Imager (Kindle Biosciences).

### Oleocanthal administration to animals

5 mg of oleocanthal was dissolved in DMSO to prepare a stock solution of 50µg/µl. The stock solution was aliquoted to avoid multiple freeze-thaw cycles, and stored at −20°C. RIP-Tag mice were intraperitoneally injected with DMSO or oleocanthal (5 mg/kg) daily starting at 9 weeks of age. Mice were weighted weekly starting from 9 weeks of age to calculate how much working solution (2.5 µg/µl) to make in normal 0.9% saline and the same dose was used for that week. Kaplan-Meier survival curve was generated using GraphPad Prism. The pancreases of treated mice at 14 weeks of age were dissected, and macroscopic tumors (> 0.5 mm^3^) were counted and measured. Tumor volume (v) was calculated using the formula for a spheroid: v = 0.52 x (width)^2^ x (length). All the tumor volumes from each mouse were summed up as the tumor burden. All procedures involving mice were approved by the Institutional Animal Care and Use Committee. There was no noticeable influence of sex on the results of this study (p value > 0.05). This study was carried out in strict accordance with the recommendations in the Guide for the Care and Use of Laboratory Animals of the National Institutes of Health. All mice were housed in accordance with institutional guidelines.

### Oil treatments

Olive oil (or Corn oil) containing treatment media was freshly prepared before each experiment by mixing oil in serum free media in a 1:25 ratio (1 mL of oil in 24 mL Media). The mixture was vigorously vortexed on highest setting for one minute on a tabletop vortex (Scientific industries Vortex Genie-2) to allow the more hydrophilic components of the oil to be extracted into the aqueous medium. The treatment media was then allowed to rest for 5 minutes and the oil settled on the top of the tube. The resulting EVOO enriched treatment media was then collected from underneath the oil layer and was used to treat the cells.

## Results

### Oleocanthal induces rapid necrotic cell death in a variety of cancer cells

As we and other groups have previously reported, oleocanthal is toxic to many cancer cells and causes rapid and extreme loss of cell viability without killing healthy cells (14, 16, 19). We treated a panel of cancer cells and normal human cells with 20 μM oleocanthal, and as expected, saw a sharp loss in viability within 24 hours among the cancer cells (MDA-MB-231 human breast cancer cells, PC3 human prostate cancer cell lines, and N134 murine PanNET cancer cells) while the non-cancerous cells (MCF10A human breast epithelial cells, HEK293T human kidney cells and BJ-hTERT human fibroblast cells) were less affected by oleocanthal treatment (Fig 1A). Phenotypic changes were observed as rapidly as one hour post treatment when cells start to round up and detach from the cell culture dishes. Moreover, loss in cell viability was induced in PC3 cells by a brief 60 min treatment of oleocanthal followed by removal of the treatment media (Fig 1B) – indicating that the cell death induced by oleocanthal is rapid. We previously reported that oleocanthal-induced cell death is due to a necrotic mechanism, whereas other groups have reported the mechanism of death to be apoptotic (15, 16).

**Fig 1.**
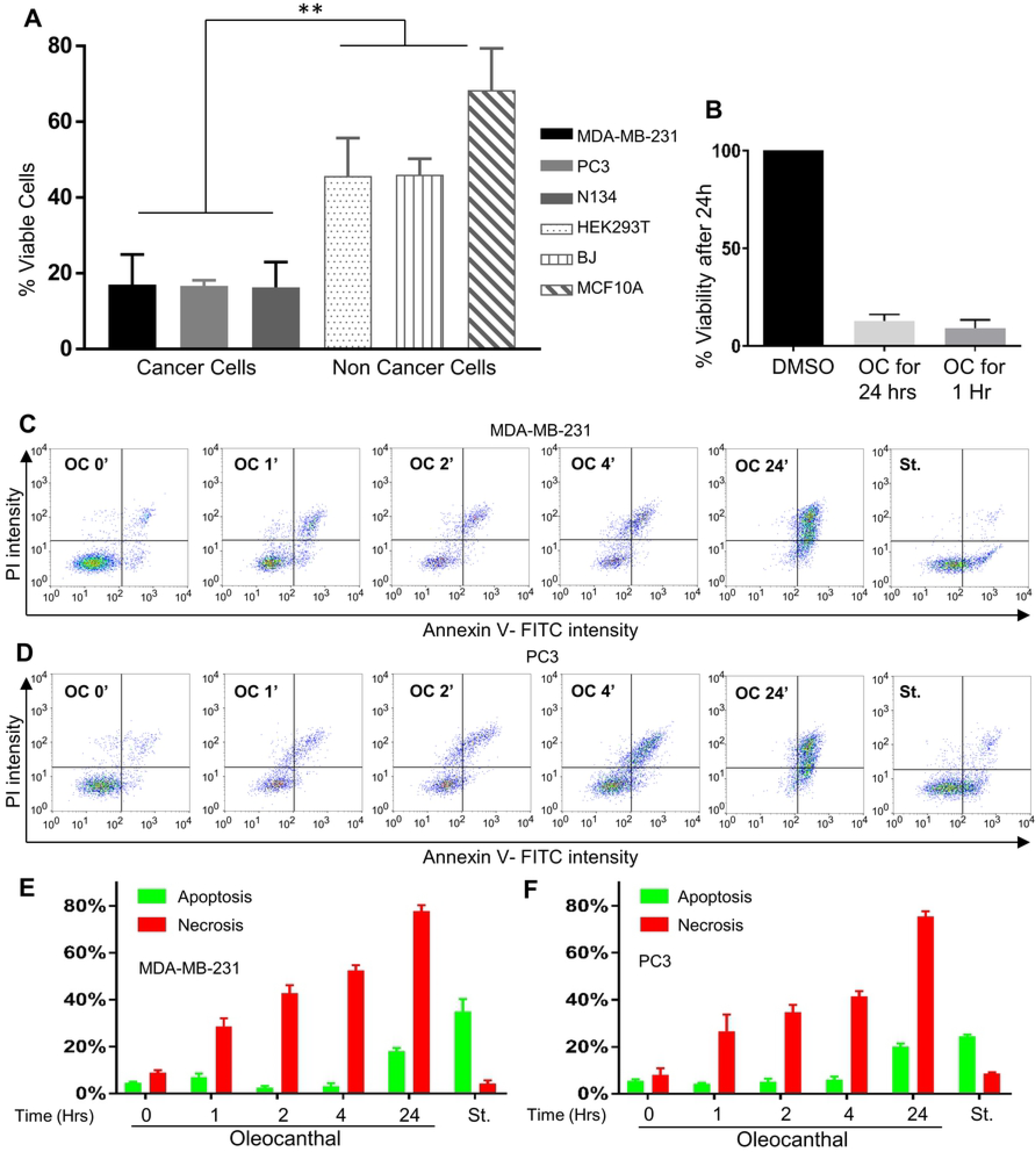
Oleocanthal induces rapid necrotic cell death in a variety of cancer cells. **(A)** The indicated cell lines were treated with 20 μM oleocanthal (OC) for 24 hours and viability was measured via the reduction of XTT. **P < 0.01 (One-way ANOVA). **(B)** PC3 cells were treated with 20μM oleocanthal or DMSO control for either 24 hours without media change, or 1 hour followed by a media change into full growth medium. Viability was measured 24 hours post treatment via the reduction of XTT. C and D) MDA-MB-231 cells **(C)** and PC3 cells **(D)** were treated with vehicle only (DMSO), or 20 μM oleocanthal for the indicated time points, and double-stained with Annexin-V FITC and PI. Fluorescence was measured on a flow cytometer (MoxiGo II). Treatment with 1μM Staurosporine (St) for 4 hours is presented as a positive control for apoptotic cells. Representative scatter plots from 3 independent experiments are shown, as well as bar graph quantifications: the lower right quadrant (apoptosis) is shown in green, and upper quadrant (necrosis) is shown in red. Bar graphs represent the mean ± SEM (n=3).

To further establish the mechanism of cell death, we performed a well-established apoptosis assay using double staining for annexin-V-FITC (AV) and propidium Iodide (PI) and compared the cell death caused by oleocanthal to the known apoptosis inducer staurosporine in MDA-MB-231 breast cancer cells (Fig 1C) and in PC3 prostate cancer cells (Fig 1D). Whereas staurosporine treated cells single-stained for AV, a hallmark of apoptosis, the oleocanthal treated cells double stained for both AV and PI – clearly distinguishing the cell death induced by oleocanthal from the apoptotic death induced by staurosporine. In the literature, double staining by PI and AV is interpreted as necrosis – although there are occasional apoptotic phenotypes observed (34). The distinction depends on whether there is an earlier time point where cells are still not permeable to PI but already stain for AV. In our hands, regardless of how short of a treatment we performed, including a 15 min treatment, we never observed oleocanthal treated cells to be single stained for AV, indicating that they do not undergo classic apoptosis. We always observed double staining for both AV and PI upon oleocanthal treatment, which led us to conclude that the mode of death was predominantly necrosis.

### Oleocanthal induces lysosomal membrane permeabilization and cathepsin release to the cytosol

In the last few years there has been a growing appreciation for the importance of lysosome-dependent cell death (35), and with this appreciation new techniques and assays have been introduced to assess LMP. The galectin translocation assay, first described by Aits et al. (32), is emerging as a gold standard to identify and quantify LMP. Galectins are β-galactoside binding proteins that normally localize to the cytosol and feature a diffuse cytosolic staining when observed in a confocal microscope. Upon damage to the lysosomal membrane, galectins translocate to damaged lysosomes and get trapped because of their affinity to luminal lysosomal β-galactoside sugars (32, 33). We performed the galectin translocation assay on MCF7 human breast cancer cells, as indicated by Aits et al., because of the high levels of galectin-3 in these cells (32, 33). We used the well-described LMP inducer, L-leucyl-L-leucine methyl ester (LLOMe) as a positive control. Within 2 hours of treatment with oleocanthal, we observed robust lysosomal staining for galectin-3, similar to LLOMe treatment (the positive control) and unlike treatment with vehicle only (DMSO) (Fig 2A). The translocation of galectin-3 from diffuse cytosolic staining to strong punctate perinuclear staining is indicative of damaged lysosomal membranes (33).

**Fig 2.**
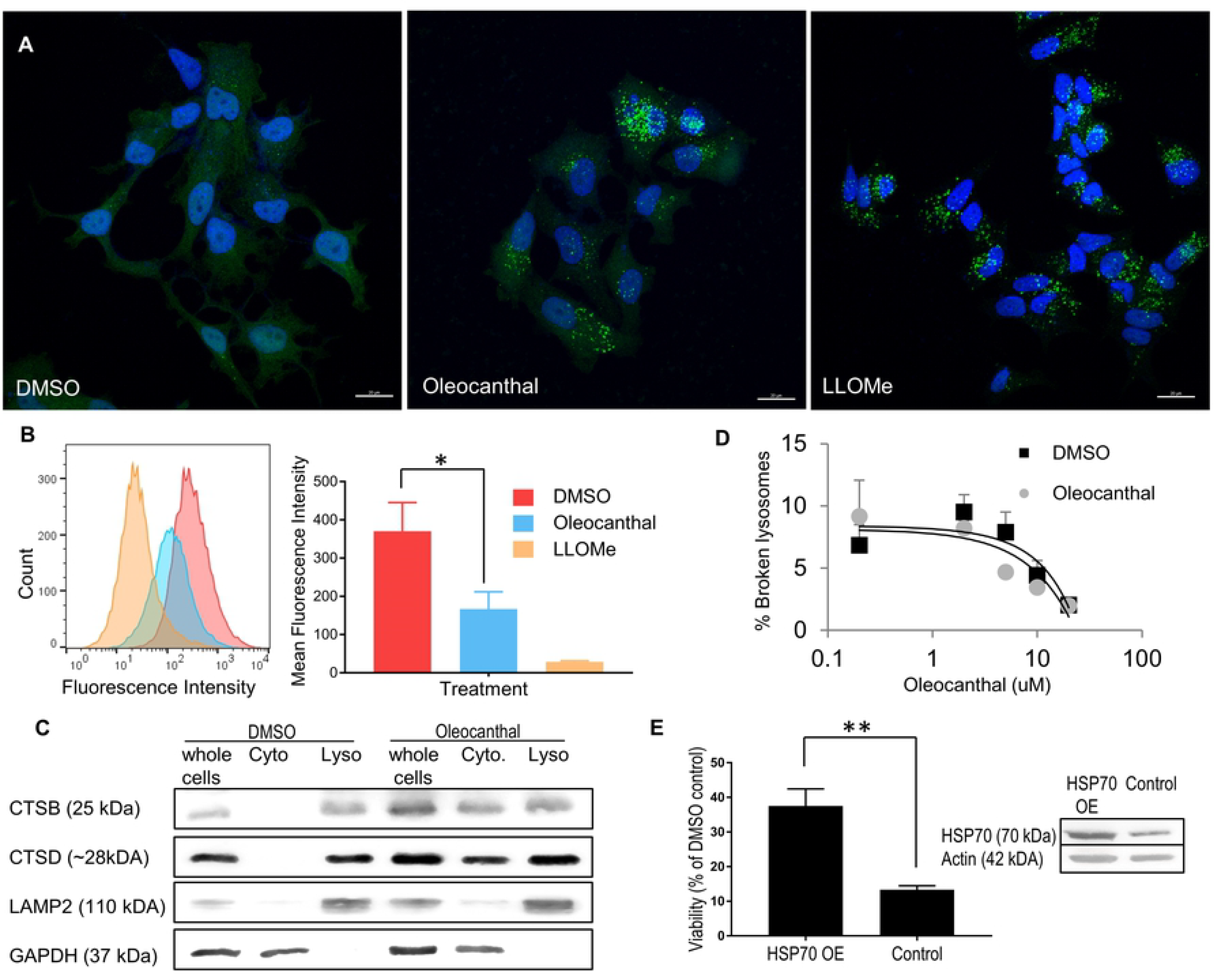
Oleocanthal induces LMP and cathepsin leakage. **(A)** MCF-7 cells were treated with DMSO, 30μM oleocanthal for 2 hours, or 2 mM LLOMe and stained for Galectin-3. Nuclei were labeled with Hoechst 33,342. Scale bars 20 μM. Green Galectin punctea indicate compromised lysosomes. **(B)** PC3 cells were treated with 20 μM oleocanthal for one hour, or 2mM LLOMe for 15 minutes, then loaded with Lysotracker green. Fluorescence intensity was measured via flow cytometry. Histogram shows a representative shift in Lysotracker fluorescence associated with perturbation to the lysosomal compartment. Bar graph shows mean fluorescence intensity of three replicate experiments. **(C)** PC3 cells were treated with 20 μM oleocanthal, and two hours later their cytosolic fractions (Cyto), and light membrane fractions containing lysosomes (Lyso) were separated. Level of cathepsin B (CTSB) and cathepsin D (CTSD) in the various fractions or whole cell lysates is shown. LAMP2 is a lysosomal marker and GAPDH is a cytosolic marker. **(D)** Lysosomes isolated from overnight serum-deprived PC3 cells were incubated for 20 min with the indicated concentrations of oleocanthal or vehicle (DMSO). At the end of the incubation, lysosomes were filtered through a vacuum manifold and b-hexosaminidase activity was measured in the flow through and in the total lysosomal fraction. Broken lysosomes were calculated as the percentage of total lysosomal hexosaminidase activity detected in the flow-through and plotted in logarithmic scale. **(E)** PC3 cells were infected with HSP70-1 Lentiviral Activation Particles, or control (scrambled) particles, and treated with 20 uM oleocanthal. Viability was assayed using reduction of XTT. *P < 0.05, **P < 0.01 (Two-tailed unpaired t-test). Bar graphs represent the mean ± SEM (n=3).

We further looked at the integrity of the lysosomal compartment by performing a LysoTracker retention assay. LysoTracker is a fluorescent acidotropic probe for labeling and tracking acidic organelles in live cells. In healthy cells, staining with LysoTracker results in a strong fluorescence signal. Loss of fluorescence is associated with either damage or de-acidification of lysosomes (36). Known LMP inducers such as LLOMe lead to decreased LysoTracker fluorescence signal within a short time post treatment (37). We, therefore, treated PC3 prostate cancer cells with oleocanthal, LLOMe, or vehicle only, and stained with LysoTracker green. oleocanthal induced a sharp reduction in fluorescence intensity (Fig 2B). Although LLOMe treated cells showed a more pronounced reduction in fluorescence intensity, oleocanthal’s effect was highly significant and further implicates LMP as the immediate cause of death in cancer cells induced by oleocanthal.

To further test whether the observed damage to lysosomes was a result of loss of acidity or actual permeability of the membrane and to assess the functional consequences of damage to lysosomes, we looked at the distribution of lysosomal enzymes in the cell. Prior to oleocanthal treatment, lysosomal hydrolases such as cathepsin B and cathepsin D were entirely excluded from the cellular cytosol (Fig 2C, 2^nd^ lane). Upon oleocanthal treatment, however, we observed a substantial release of these proteases to the cytosol (Fig 2C, 5^th^ lane), indicating that oleocanthal treatment causes cathepsins to be released from the lysosomes to the cytosol.

Interestingly, incubating purified lysosomes isolated from PC3 cells *in vitro* with increasing concentrations of oleocanthal (as we don’t know the final cytosolic concentration of oleocanthal inside cells) had no appreciable difference in lysosomal stability as compared to vehicle (Fig 2D). This indicates that oleocanthal does not act directly as a membrane disrupting agent on lysosomes, but rather induces lysosomal permeability only in a cellular context – likely through oleocanthal metabolites.

The heat shock protein HSP70 is known to stabilize lysosomal membranes (28) and in various models of LMP, HSP70 provides protection from subsequent cell death (38). We, therefore, overexpressed HSP70 in PC3 prostate cancer cells and examined the effect on oleocanthal-induced loss of cell viability. Indeed, oleocanthal-induced loss of cell viability was partially rescued by HSP70 overexpression (Fig 2*E*), further supporting a role for LMP as the cause of oleocanthal-induced cell viability.

Collectively the data provided in Fig 2 strongly suggest that oleocanthal triggers rapid damage to lysosomes, which causes them to become permeable and leaky, allowing cytosolic proteins into the lysosome (galectin-3) and lysosomal proteins (cathepsins) out into the cytosol. The rapid assault on lysosomes, on which cancer cells are highly metabolically dependent supports the idea that the cellular toxicity caused by oleocanthal is due to LMP. All other observed effects of oleocanthal on apoptotic and necrotic forms of cell death in cancer cells are likely down-stream of the LMP and dependent on the corresponding degree of LMP.

### Oleocanthal extends the life span of mice bearing PanNET tumors

To assess the benefit of oleocanthal treatment in a genetically engineered mouse model of PanNET, *RIP-Tag* mice (39), we performed a mouse survival trial. The *RIP-Tag* mice inevitably develop tumors that progress through well-defined stages that closely mimic those found in human pancreatic neuroendocrine tumorigenesis (i.e., hyperplasia, angiogenesis, adenoma, and invasive carcinoma) (39). We treated mice with 5 mg/kg oleocanthal or DMSO vehicle daily through intraperitoneal injection starting at 9 weeks of age. The median survival of vehicle-treated mice was 14 weeks of age (Fig 3A). In contrast, the oleocanthal-treated animals had a significant extension of life surviving a median period of 18 weeks – or an additional 4 weeks. To determine the effect of oleocanthal on tumor sizes, we treated another cohort of mice with 5 mg/kg oleocanthal or vehicle DMSO daily through intraperitoneal injection starting at 9 weeks of age and euthanized them at 14 weeks of age (5 week treatment). Although the effect on tumor burden did not reach statistical significance, there was a trend toward smaller tumor burden with oleocanthal treatment (Oleocanthal: 14.7 mm^3^ vs. DMSO: 24.8 mm^3^) (Fig 3B).

**Fig 3.**
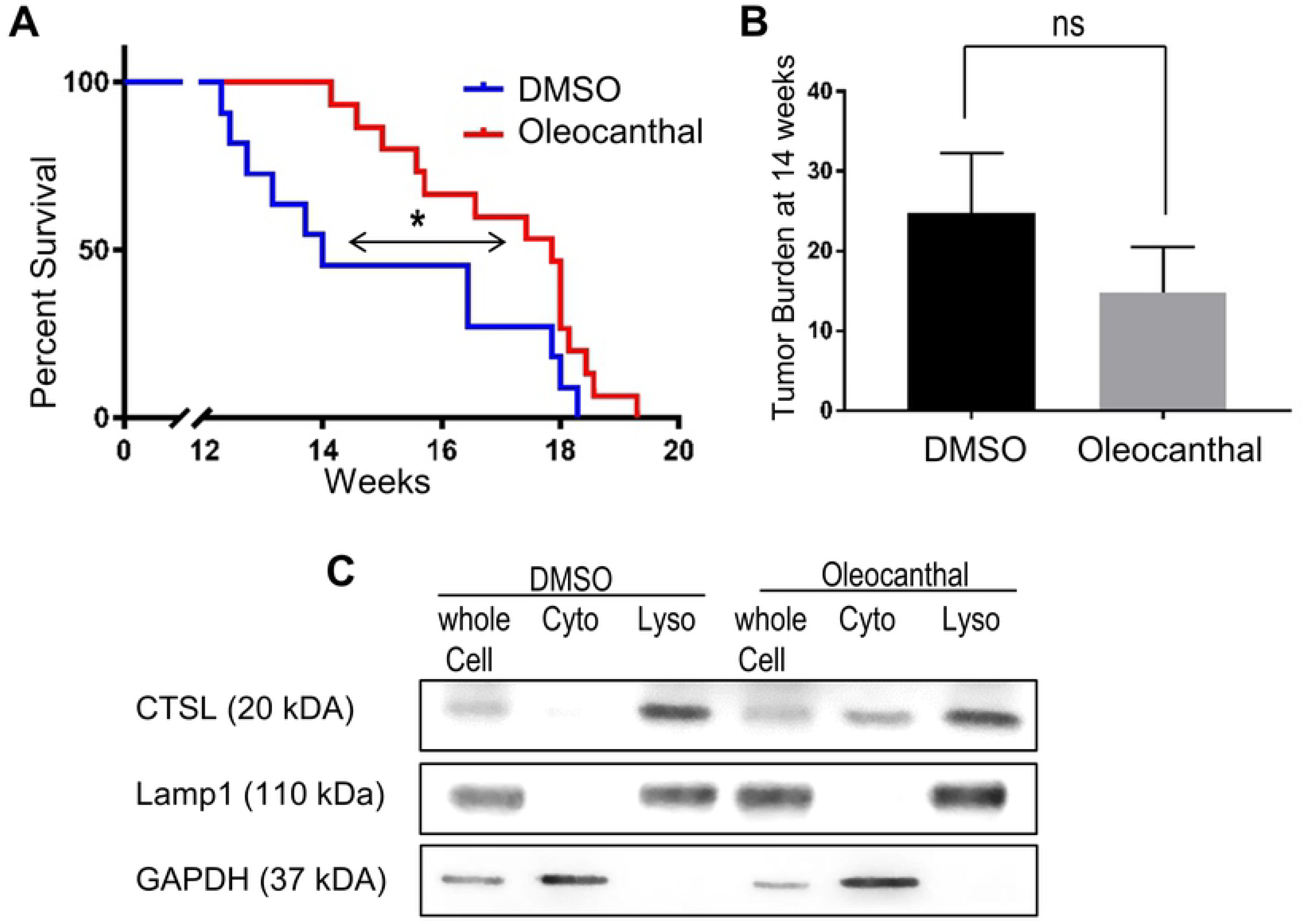
Oleocanthal increases life span of mice with PanNET tumors. **(A)** Kaplan-Meier survival curve for RIP-Tag mice receiving DMSO or oleocanthal. The mice were treated with DMSO (n= 11) or oleocanthal (5 mg/kg, n=15), 7 days a week. Mice were treated starting from 9 weeks of age. Both the Gehan-Breslow-Wilcoxon method and the Log-rank (Mantel-Cox) method were used to calculate statistical significance *P < 0.05. **(B)** Tumor burden from mice treated with DMSO or oleocanthal (n =7 for each group) starting from 9 weeks of age and ending at 14 weeks of age. ns P > 0.05. **(C)** A cell-line derived from a murine PanNET tumor, was established (N134). Cells were treated with DMSO or oleocanthal and analyzed for cytosolic cathepsin L (CTSL) via Western blot as in Fig 2C.

To determine whether the tumors were smaller due to LMP induced cell death, we checked for cytosolic cathepsin release in the murine cells. We treated an established cell line derived from a *Rip-Tag* neuroendocrine tumor (N134) *in vitro* and found cathepsin L (which is a highly expressed cathepsin in N134 cells) present in the cytosol upon oleocanthal treatment (Fig 3C). The data presented in Fig 3 provide evidence that oleocanthal suppresses tumorigenesis in a mouse model for PanNET pancreatic neuroendocrine tumors.

### Oleocanthal-rich EVOOs are toxic to cancer cells via LMP

The use of EVOO in the Mediterranean diet has been associated with cancer protective effects (4). However, the concentration of oleocanthal in EVOOs varies greatly (31). We, therefore, examined the effect of EVOOs with varying oleocanthal concentrations on cancer cell viability. We hypothesized that EVOOs with high levels of oleocanthal will show greater toxicity towards cancer cells than EVOOs with lower levels of oleocanthal. The levels of oleocanthal present in several EVOOs, a non-virgin olive oil, and corn oil were determined by 1H-NMR as described in Materials and Methods (Fig 4A). Two EVOOs (Colavita EVOO, and Olive Ranch) had average content of oleocanthal. Two EVOOs (The Governor and Atsas) had levels of oleocanthal that was 5 or 6-fold higher than the other EVOOs. The non-virgin olive oil (Colavita mild) and the corn oil (Mazola) had no detectable oleocanthal and were used as negative controls. We then prepared cellular treatment media that consisted of cell culture media and EVOO in a ratio of 25:1. We used this specific ratio because it would make the maximum oleocanthal level in the treatment media in the 20 μM range for the most potent EVOO. To ensure that the oleocanthal was transferred to the media, we vortexed the mixture vigorously, in essence extracting the more polar components (the phenolic content of the oil) into the media. We then treated PC3 prostate cancer cells (Fig 4B) and MDA-MB-231 breast cancer cells (Fig 4D) with this enriched media. Strikingly, the ability of the EVOO enriched media to kill the cells was directly and linearly correlated to the EVOO’s oleocanthal content. The oils with the highest oleocanthal content reduced cell viability for both PC3 and MDA-MB-231 cells to a similar degree to that observed in response to purified oleocanthal. The oils with the next two highest oleocanthal concentrations reduced viability in a manner corresponding with oleocanthal concentration; and the oils with no measurable amounts of oleocanthal did not affect cell viability relative to the no-oil negative control treatment. We also analyzed the ability of EVOO to induce LMP as determined by cathepsin release. As shown in Figs 4C and 4E, the EVOOs with the highest concentration of oleocanthal induced cathepsin release caused leakage of both cathepsin D and B into the cytosol of PC3 cells (Fig 4C) and MDA-MB-231 cells (Fig 4E). In contrast, the other oils caused minimal cytosolic cathepsin release - indicating that the oleocanthal content in EVOOs is a major determinant for EVOO’s cancer-protective properties. These data demonstrate that oleocanthal is able to exert this beneficial effect when delivered via whole EVOO and not only in a purified phenolic form.

**Fig 4.**
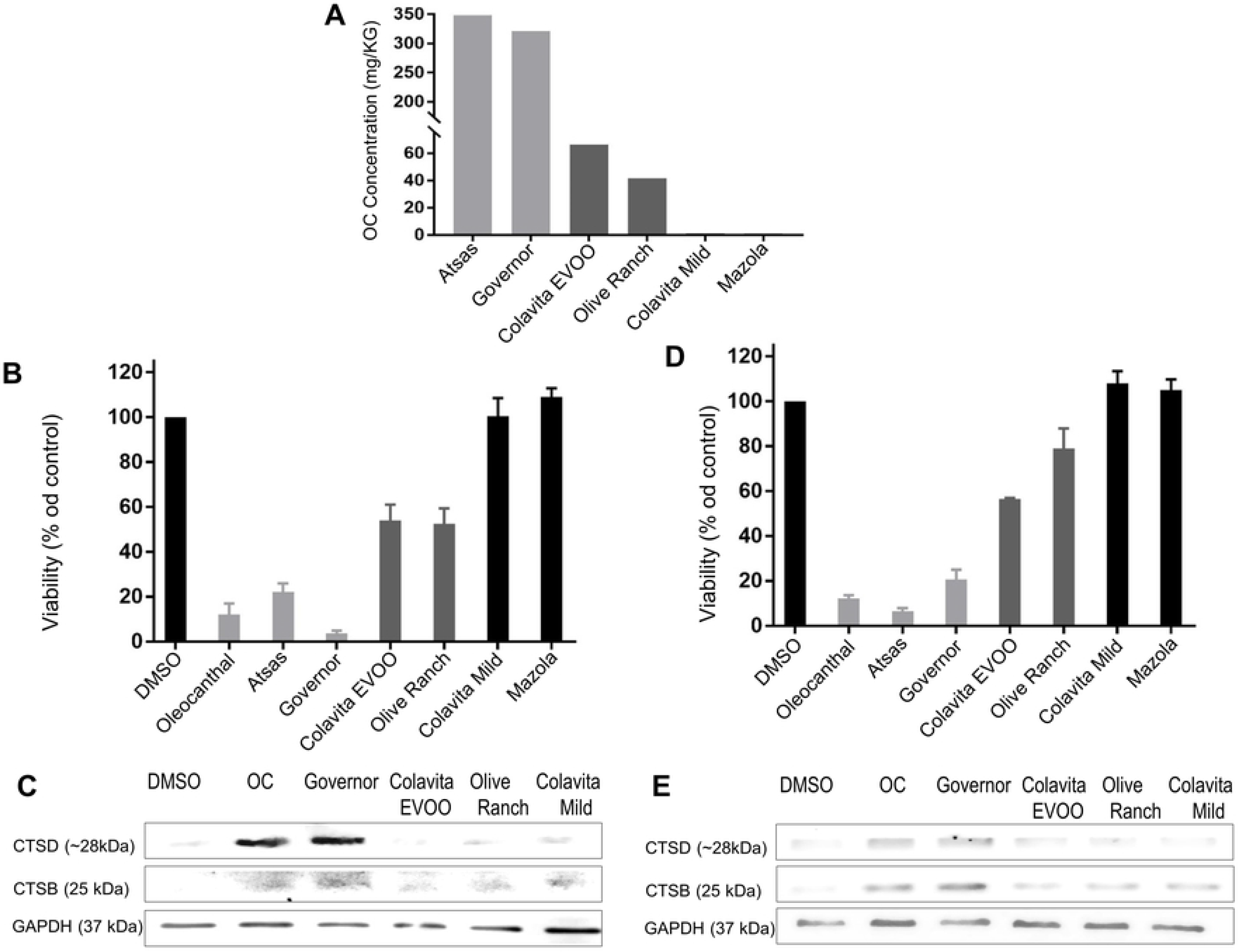
Oleocanthal-rich olive oils are toxic to cancer cells via LMP. **(A)** Relative oleocanthal concentration in various oils was measured by H1 NMR as described in Materials and Methods. **(B and C)** PC3 cells **(B)** and MDA-MB-231 cells **(C)** were treated with 20 μM OC, or the specified oils for 24 hours. Viability was measured via the reduction of XTT. **(D and E)** Cytosolic lysates were collected as in Fig 2C and subjected to Western blot analysis of cathepsin B (CTSB) and cathepsin D (CTSD) in the cytosol. Bar graphs represent the mean ± SEM (n=3)

## Discussion

Although several groups have demonstrated oleocanthal’s ability to inhibit key proteins that promote cell growth and survival (12, 14-17, 40), a unifying mechanism for the specific and irreversible cellular death-inducing properties of oleocanthal has not been established. In this report, we observed that a transient exposure of cancer cells to oleocanthal for one hour resulted in the loss of cell viability after 24 hours. Although a classic apoptotic mechanism has been proposed (16, 20, 21), in our hands the rapid cell death caused by oleocanthal was necrotic. Specifically, viable cells were not observed to display phosphatidylserine on the outer membrane leaflet as evidenced by staining with AV, a well-established phase in the apoptotic cascade. Furthermore, using three different and complementary methods, we demonstrated that oleocanthal-treated cells undergo LMP. The latest, most robust method to assess LMP is the galectin translocation assay (33). We observed that oleocanthal treated MCF7 breast cancer cells showed robust galectin-3 translocation to lysosomes, similar to that observed with the established LMP inducer LLOMe. In a biochemical assay that checks the leakage of lysosomal enzymes into the cytosol, we observed a pronounced leakage of both cathepsin D and cathepsin B to the cytosol in PC3 prostate and MDA-MB-231 breast cancer cells. The translocation of cathepsins of two different sizes suggests that the lysosomal membrane undergoes severe and unrepairable permeabilization. Agents that are known to cause LMP with only minimal cathepsin release, such as LLMOe (37) enable cells to survive the initial LMP and repair their lysosomal membrane. Other agents that cause the release of cathepsin D (a small hydrolase) but not the release of cathepsin B (a larger hydrolase) are often associated with apoptosis (22). We, therefore, conclude that the degree of lysosomal damage in the case of oleocanthal is massive and leads to rapid necrosis in the affected cancer cells with less and survivable damage to normal cells.

It was previously suggested that many cancer cells are more vulnerable to attacks on their lysosomes because they have larger and more fragile lysosomes (41) and are more reliant on lysosomal processes metabolically (27). Furthermore, many cancer cells upregulate lysosomal biogenesis and lysosomal enzyme turnover (27). Therefore, once lysosomal enzymes and acids are released into the cytosol en mass, rapid cell toxicity ensues (42). The effect of oleocanthal was observed in both cell culture and a live mouse model for the development of PanNETs (39) where lifespan was extended by 4 weeks (29%). It has been reported that 2.6 adult mice days are equivalent to one human year (43). Based on this life-span conversion, oleocanthal might extend life 10.4 years for PanNET pancreatic neuroendocrine cancer patients. Importantly, the cancer cells from the PanNETs when put in culture released cathepsin upon oleocanthal treatment and died rapidly.

In addition to looking into the effects of purified oleocanthal, we were very interested to see if oleocanthal in a more natural form can cause a similar outcome. Since different olive oils are known to have varied oleocanthal concentrations as a function of their origin, harvest time, and processing methods (7), we examined several olive oils with varied concentrations of oleocanthal from very low to very high. For our *in vitro* experiments, we used two EVOOs with average low oleocanthal content and two with very high oleocanthal content (about 5 times the average), and for our negative control we used two oils that contained no measurable oleocanthal. Upon treatment of cultured cancer cells with oil enriched cell culture media we observed that the concentration of oleocanthal in the oil was directly related to the toxicity of the oils towards cancer cells. The oils with the high oleocanthal content completely killed the cancer cells in a manner similar to purified oleocanthal. The oils with the average oleocanthal content, also reduced viability but to a lesser extent. The non EVOOs with no oleocanthal had no effect on cell viability. Furthermore, by looking at cytosolic cathepsin release, the EVOOs mechanism of promoting cancer cell death also involved LMP, similar to the effects of purified oleocanthal.

Many studies have linked consumption of EVOO with reduced incidence of cancer (4), most significantly a randomized trial in which elevated EVOO in the diet led to a 62% reduction in the incidence of breast cancer in Spain over a 5 year period (5). Data provided here link the cytotoxic effects of EVOOs to their level of oleocanthal. The cytotoxic effects were due to the ability of oleocanthal to induce LMP and necrotic cell death preferentially in cancer cells. Whereas pure oleocanthal can also have negative effects on non-cancerous cells, EVOO is considered safe and healthy and, therefore, could be both preventative as well as a potential treatment – as indicated by the Spanish study (5). Since the apparent target for oleocanthal-induced necrosis is the lysosome, the reason for the elevated sensitivity of cancer cells to oleocanthal could be due to the increased size and fragility of the lysosomal compartment of cancer cells (27). If the enlarged fragile lysosomal compartment (23, 41) is the reason for increased sensitivity to oleocanthal, it is likely that EVOOs with high oleocanthal could be preventative for many cancers – in addition to reduced breast cancer in Spain (5). Whether purified oleocanthal could be used therapeutically remains to be evaluated. How can one determine whether there are high levels of oleocanthal in an EVOO? EVOOs with high oleocanthal levels produce a unique stinging sensation in the back of the throat and not elsewhere in the mouth, as well as eliciting a brief coughing that has been used to determine the presence of oleocanthal in EVOO (8). Tasting EVOO for this signature stinging sensation and cough elicitation could allow people to identify EVOOs with high oleocanthal content without sophisticated equipment. In light of the results presented in this report, and since EVOOs have been safely used in the diet for millennia and are associated with good health, the authors believe that consuming more EVOO with high oleocanthal content is a prudent dietary approach to cancer prevention with the caveat that dietary oils convey calories and consequently other caloric sources will have to yield to avoid obesity.

## Acknowledgments

We thank Dr. Ana Maria Cuervo for valuable guidance, and to Daniela Mikhaylov, Lucy Pascarosa, and Ismat Zerin for experimental assistance.

